# Design, construction and characterisation of a novel nanovibrational bioreactor and cultureware for osteogenesis

**DOI:** 10.1101/543660

**Authors:** Paul Campsie, Peter G. Childs, Shaun N. Robertson, Kenny Cameron, James Hough, Manuel Salmerón-Sánchez, Monica P. Tsimbouri, Parag Vichare, Matthew J. Dalby, Stuart Reid

## Abstract

In regenerative medicine, techniques which control stem cell lineage commitment are a rapidly expanding field of interest. Recently, nanoscale mechanical stimulation of mesenchymal stem cells (MSCs) has been shown to activate mechanotransduction pathways stimulating osteogenesis in 2D and 3D culture. This has the potential to revolutionise bone graft procedures by creating cellular graft material from autologous or allogeneic sources of MSCs without using chemical induction. With the increased interest in mechanical stimulation of cells and huge potential for clinical use, it is apparent that researchers and clinicians require a scalable bioreactor system that provides consistently reproducible results with a simple turnkey approach. A novel bioreactor system is presented that consists of: a bioreactor vibration plate, calibrated and optimised for nanometre vibrations at 1 kHz, a power supply unit, which supplies a 1 kHz sine wave signal necessary to generate approximately 30 nm of vibration amplitude, and custom 6-well cultureware with toroidal shaped magnets incorporated in the base of each well for conformal attachment to the bioreactor’s magnetic vibration plate. The cultureware and vibration plate were designed using finite element analysis to determine the modal and harmonic responses, and validated by interferometric measurement. This helps ensure that the vibration plate and cultureware, and thus collagen and MSCs, all move as a rigid body, avoiding large deformations close to the resonant frequency of the vibration plate and vibration damping beyond the resonance. Assessment of osteogenic protein expression was performed to confirm differentiation of MSCs after initial biological experiments with the system, as well as atomic force microscopy of the 3D gel constructs during vibrational stimulation to verify that strain hardening of the gel did not occur. This shows that cell differentiation was the result of the nanovibrational stimulation provided by the bioreactor alone, and that other cell differentiating factors, such as stiffening of the collagen gel, did not contribute.

## Introduction

Increases in life expectancy, or lifespan, seen throughout the developed world are a valuable indicator for the progress of modern medicine. However, a less quantifiable measure is quality of life during those additional years; this is termed healthspan. The incidence of skeletal injuries due to age related conditions such as osteoporosis and osteoarthritis provide one such metric. As such, development of treatments which lead to increased bone density or fracture healing are prime targets for the regenerative potential of mesenchymal stem cells (MSCs)^1, 2^.

The controlled osteogenesis of MSCs through mechanical stimulation has been demonstrated through several methods including passive and active strategies. Passive methods such as altered substrate topography and environmental stiffness provide one mechanism based on altering the adhesion profile^3–7^, whilst active methods include exposure to variations of force from external sources^8–13^. Centrifugation, vibration and shear flow have all provided increases in osteogenesis through external modulation of the force imposed on the cell structure. The use of vibration as a mechanotransductive stimulus has been explored with varied vibrational parameters^14–16^. Vibration of periodontal ligament stem cells at 50 Hz with a peak acceleration of 0.3 *g* showed increased markers of osteogenesis^17^ whilst another study of adipose-derived stem cells stimulated using a feedback controlled vibration source at 50 and 100 Hz with a reported peak acceleration of 3 *g*, showed increased levels of alkaline phosphatase (ALP) activity and mineral deposition, however not at the same level produced by osteogenic media^18^.

Initial studies from Curtis *et al.*^19^ and Nikukar *et al.*^20^ conducted using vibration amplitudes of tens of nanometres showed that endothelial cells and MSCs are sensitive to vibrations of this level. To achieve nanometre vibrations, several studies^19–22^ used a single piezo actuator, single Petri dish apparatus to produce accurate vertical vibration over a small growth surface, however this imposed limitations of scale and vibration amplitude which become relevant if developing towards a larger industrial process. The use of alternative growth containers (*e.g.* T-75 flasks) and cultureware, along with a larger amplitude range (and thus cellular forces), requires a more uniform, versatile and reusable bioreactor platform. An initial prototype bioreactor was constructed incorporating an array of piezos attached to a metal platform for multi-well cultureware and is detailed in a study by Tsimbouri *et al.*^23^ addressing both of these issues. The aim of the work presented in this article is to progress this design towards a Good Manufacturing Practice (GMP) compatible system which is suitable for the purpose of supplying a small scale clinical trial.

The proposed bioreactor system consists of three main elements: a bioreactor vibration plate, a power supply to drive the piezo array underneath the vibration plate, and custom 6-well cultureware that can magnetically attach to the vibration plate. Finite element analysis (FEA) was used to improve the bioreactor design as well as optimising its construction, minimising large amplitude variance arising from resonances or substrate deformation. These factors are important when using nanoscale amplitudes as consistent vibration ensures that all cells on the bioreactor experience the same levels of acceleration. Previous experiments^19–23^ used vibration amplitudes of 15 – 35 nm and frequencies of 1 Hz – 5 kHz and the most significant osteogenic effects were observed at 1 kHz^20, 22^. It is important that the bioreactor operates within this frequency range and crucial that this range is well below the bioreactor’s resonant condition to avoid resonant amplification/damping. FEA was also used in the design of the cultureware to confirm that the base of each well would give a uniform displacement when vibrated at 1 kHz and guarantee that resonant modes would not arise in the experimental operating window. Once constructed, laser interferometry was utilised to accurately measure the vibration displacement from the bioreactor top plate and the wells of the cultureware, thus validating the FEA models. The power supply was designed to output a pure sine wave at 1 kHz with an amplitude of 30 nm using a direct digital synthesis (DDS) waveform generator and a parallel configuration of power amplifiers. Using a reconstruction filter to remove high frequency components of the DDS output made certain that the 1 kHz sine wave was as pure as possible.

Finally, the overall operation of the bioreactor system was validated through biological experiment by quantification of osteogenic protein expression of MSC’s exposed to nanovibrational stimulation. Atomic force microscopy (AFM) measurements were also carried out on collagen gel used in these experiments to determine that the vibrations were transmitting from the cultureware into the gel and that the stiffness of the gel didn’t significantly increase while being nanovibrated through non-Newtonian/strain hardening effects^24^.

## Results

### Design of bioreactor vibration plate using FEA

The bioreactor construction, particularly material choices, piezo positioning, and cultureware attachment, was designed to optimise the delivery of nanoscale vibration across a frequency band relevant to the previous nanovibrational studies, between 1 Hz and 5 kHz. The general approach was to ensure that the resonant conditions of the top plate and base were suitably above the frequency of operation, to prevent resonant amplification/damping. With this condition met, the top plate should move as a rigid body, providing consistent and quantifiable amplitudes of vibration.

The basic design of the bioreactor vibration plate is an array of piezos sandwiched between a heavy aluminium base block and a lighter bimetallic top plate (comprised of stainless steel and aluminium). The aluminium side of the top plate faces the piezo array and base block which allows cultureware to be magnetically attached. The heavy base block provides inertial mass ensuring that the majority of the piezo expansion is directed towards the relatively lighter top plate while the bimetallic top plate allows its weight to be kept as low as possible (rather than using a single stainless steel plate), maximising the resulting piezo displacement, and keeping the fundamental resonant mode of the plate above the bioreactor operating condition. To determine appropriate dimensions of the top plate that would result in a 1st order resonant mode above the operating frequency range, FEA was performed in ANSYS Workbench 17.1 (ANSYS Inc, Pennsylvania, USA) to evaluate the modal and harmonic response of various designs. For all models in this article that include the top plate, a displacement of 30 nm is applied at the piezo contact points on the aluminium side of the top plate in order to determine the transmission through the entire structure. The opposite surface of the piezos (not in contact with the aluminium plate) are fixed points. The modal analysis showed that a top plate with a length of 176 mm and width of 128 mm (which could comfortably accommodate 2 standard 6-well cultureplates), and comprised of 5 mm thick aluminium 8092 and 5 mm thick stainless steel 420 bonded with a 0.1 mm layer of Araldite 2012 (Araldite, Basel, Switzerland), has a fundamental resonant mode at 8230 Hz. Therefore, biological experiments will not be adversely affected by the greatly increased displacement at this resonance.

To create the bioreactors in a cost-effective manner it was estimated that thirteen to fifteen piezos would be optimum, with the aim of 10% variation in displacements with the minimum number of piezos. Harmonic response FEA modelling of a thirteen piezo array, in alternate rows of three and two, and a fifteen piezo array, in rows of three, showed that the thirteen piezo array creates the most uniform displacement across the top plate (Figure 1). In the fifteen piezo array all actuators are aligned, creating alternating lines of piezo anchored top plate and free-floating top plate. This produces distinct alternating bands of minimum and maximum displacement which would lead to cells receiving inconsistent levels of vibration across the attached cultureware. The thirteen piezo array has the actuators arranged in a checkerboard layout avoiding distinct regions of low/high vibration amplitude. Larger variations in displacement amplitude are mainly confined to the edges which are not direct regions of contact with attached multi-well plates.

**Figure 1.**
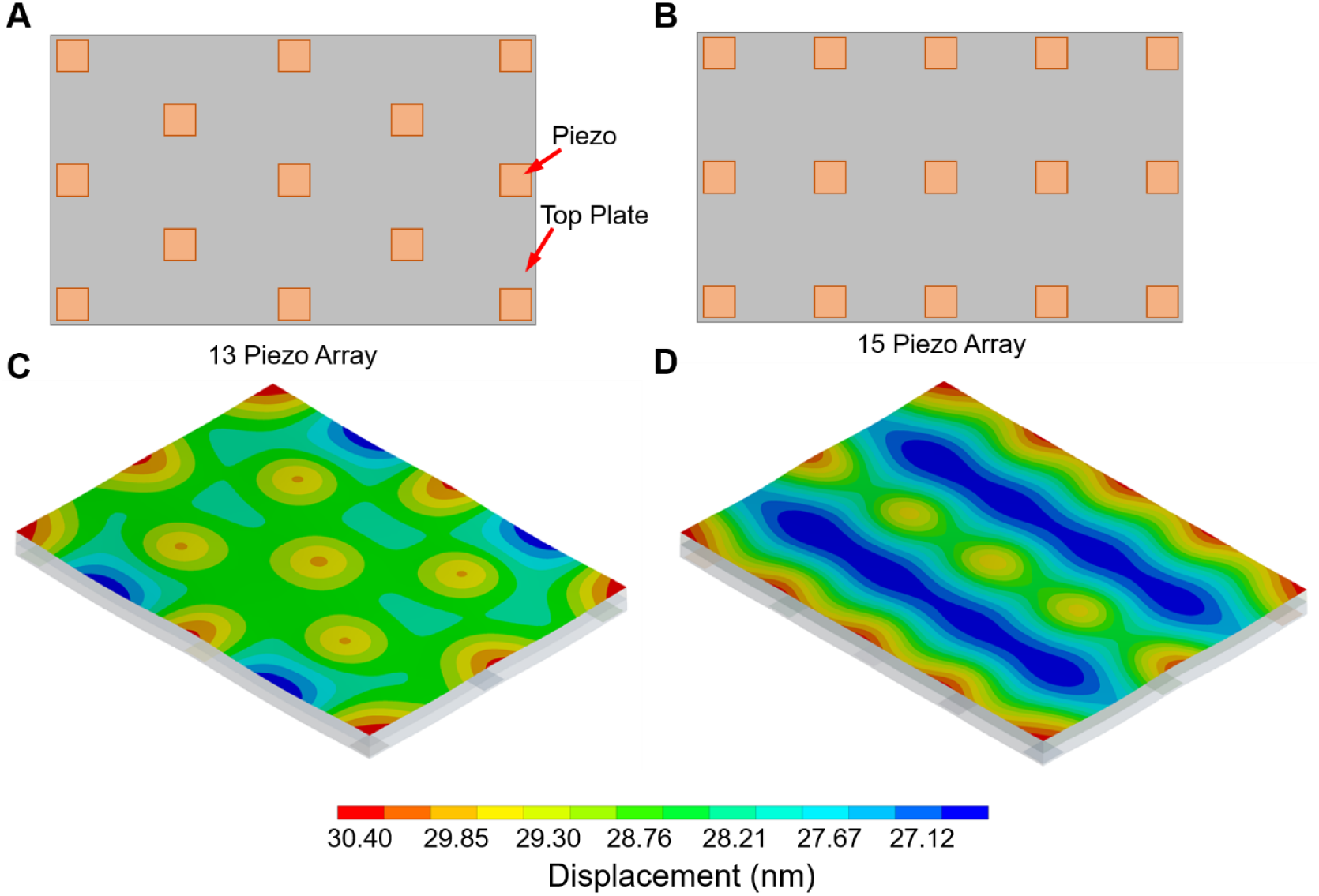
FEA analysis was performed in ANSYS workbench 17.1 to determine the harmonic response at 1 kHz on the thirteen and fifteen piezo array top plate arrangement. (A) Diagram of thirteen piezo array (B) Diagram of fifteen piezo array (C) Predicted nanoscale displacement of thirteen piezo array at 1 kHz (D) Predicted nanoscale displacement of fifteen piezo array at 1 kHz.

Aside from the structural resonances of the bioreactor, the piezo actuators have an intrinsic resonant frequency (*f*_0_). For a piezo mass (*m*) and given loading mass (*M*) the actual resonant frequency 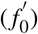 varies as Equation 1^25^.

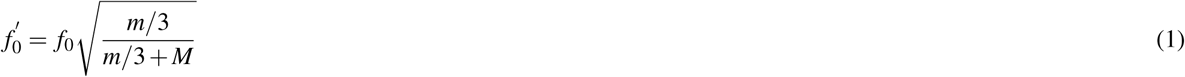

For our particular set-up, it is assumed that *M* is the total mass of the complete top plate, plus two 6-well plates with 4 ml of media in each well, equally spread over the thirteen piezos giving a loading mass of approximately 107 g per actuator. With an actuator mass (*m*) of 1.56 g this results in an estimated resonance of 41.8 kHz which is far outside the operating frequency range for the experiments reported here. It was found that a 1 ml change in media in each well results in a change in resonant frequency of approximately 100 Hz. For example, 1 ml of media in each well produces a resonant frequency of 42.1 kHz and filling each well with 7 ml of media gives a resonant frequency of 41.5 kHz.

### Custom cultureware design

Previous experiments^23^ have used standard multi-well cultureware with either circular ferrite magnets or a sheet of rubber magnet bonded to the bottom surface. Even though this is a practical solution, there is still the possibility for variation between experiments depending on exactly where/how the magnets are bonded. In addition, standard cultureware has a rigid, inflexible, base which can increase the possibility of insufficient contact, resulting in poor vibration transmission, if not aligned precisely with the vibration plate. This highlights the need for bespoke cultureware that integrates well with the bioreactor, requires no involvement from the end user to setup, and can also be produced on an industrial scale. This led to the design of an insert type, multi-ejection stroke injection mould for producing 6-well plate cultureware with toroidal shaped, unidirectional ferrite magnets embedded in the base of each well.

The injection moulded, polyproylene (PP), cultureware presented here features a more flexible design to those commercially available through the use of ‘floating’ wells with only two points of contact with the plate frame. This allows the wells to be in rigid magnetic contact with the bioreactor top plate while the frame remains flexible. In addition, it also allows individual clinical samples to be detached from the frame, sealed and easily transported/stored. Halbach array magnets were used because they only have a magnetic field on one face while the magnetic field on the opposing face is reduced to almost zero. This prevents any significant magnetic field interacting with the cells (multi-tesla magnetic fields can lead to DNA alteration^26^) while still providing appropriate coupling between the bottom surface of the wells of the cultureware and the bioreactor top plate. The toroidal shape of the magnets allows optical viewing of the cells during experiments. Details on the injection mould tooling can be found in Supplementary Results and Methods online.

As with most polymer materials, modification of the surface chemistry is required to aid cell adhesion and proliferation due to lower surface energies^27^, which can be increased via plasma surface activation^28, 29^. Air-based plasma treatment (Plasma cleaner, Harrick Plasma, New York, USA) was applied to the PP cultureware for different time durations (from 0 – 7.5 mins) to find a minimum level of plasma treatment required to sufficiently alter the surface chemistry from hydrophobic to hydrophilic, promoting the adhesion and proliferation of human osteoblast-like MG63 cells. An indicator of the surface energy of a polymer is its wettability^30^ which can be determined by measurements of its water contact angle (WCA). Post-plasma treatment the WCA on the PP cultureware was monitored over a period of one week and it was found that plasma treatment for at least 5 minutes is required to significantly alter the WCA, allowing the adhesion of MG63 cells (Figure 2).

**Figure 2.**
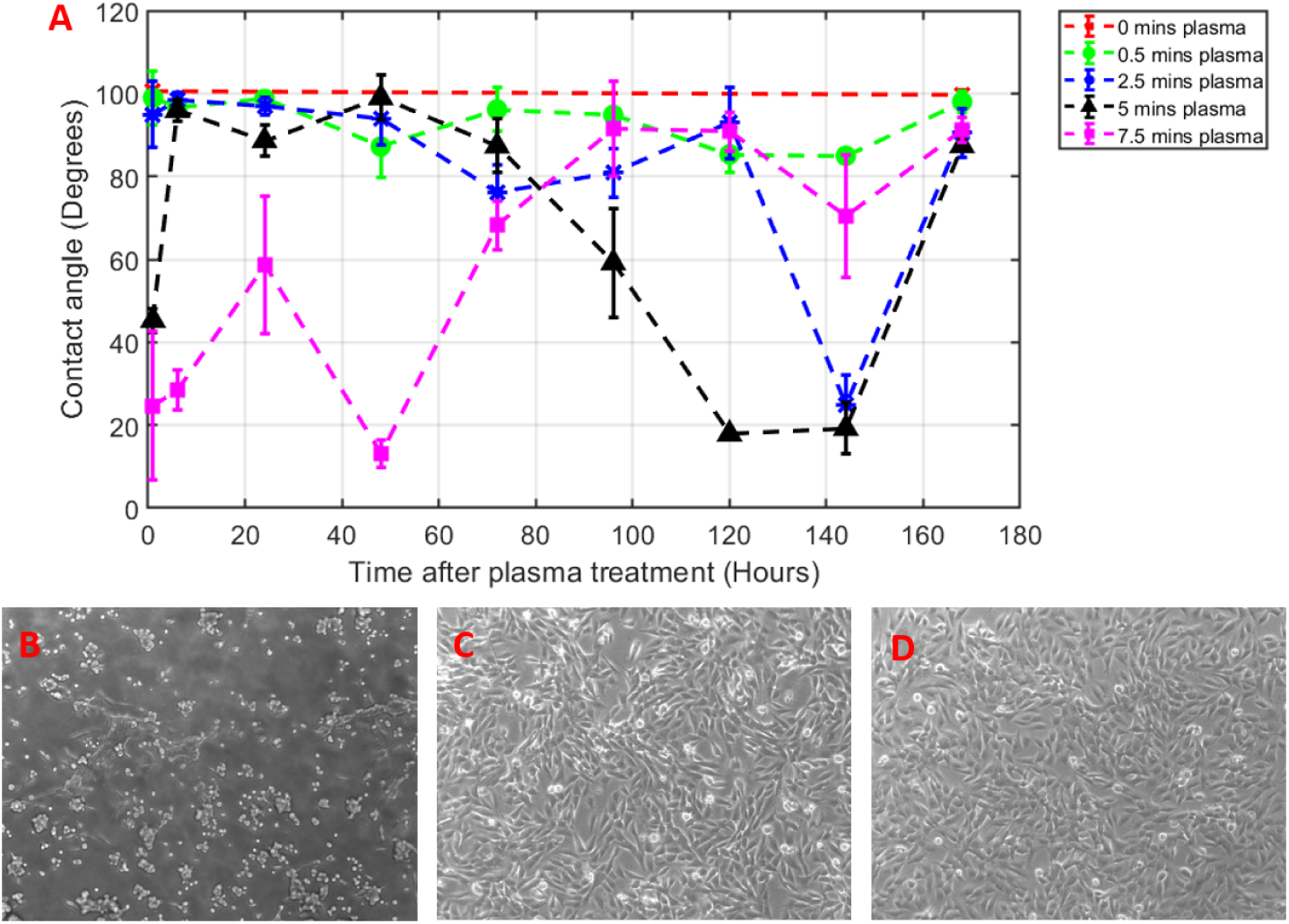
Water contact angle measurements of PP cultureware after different doses of plasma treatment and microscopy images of MG63 cells on PP and polystyrene (PS) 6-well plates. A plot of WCA measurements post plasma treatment (A) shows that at least 5 minutes is required to significantly alter the WCA to a level that would allow cells to adhere and proliferate. Images of (B) non-adherence of MG63 cells on the PP 6-well plate prior to plasma treatment, (C) adhesion and proliferation of MG63 cells on plasma treated PP 6 well plate, and (D) MG63 cells cultured on a standard Corning PS 6-well plate.

These measurements have demonstrated proof of principle that plasma activation of the PP cultureware can provide drastic improvements on WCA and favourable cell adhesion, however, the process will require further development to ensure stability and longevity of shelf life. Further study would also be required to find the optimum WCA for cell adhesion on this PP cultureware, however, an initial estimate could be in the range 60^*◦*^ – 80^*◦*^ based on studies involving MG63 cells on PS^30^ and L cells on PP^31, 32^.

FEA was performed on the 6-well plate design with a modal analysis revealing that the cultureware has a fundamental mode at 379 Hz and some modes close to 1 kHz (for table of modes 1 to 12 see Supplementary Table S1). Even though this is within the frequency range used for biological experiments with the bioreactor, the analysis shows that the wells themselves are unaffected, with largescale displacements confined to the frame of the 6-well plate, as shown in Figure 3. A resonance mode that would directly affect the bottom surface of the wells in the frequency range 1 Hz to 5 kHz could not be found in our analysis. Some modes in this range were found to affect the sides of the wells, however, the strict vertical vibration applied by the bioreactor is unlikely to significantly excite these particular modes. To understand how the displacement propagates through the top plate and into the wells of the cultureware, a harmonic analysis was performed with the 6-well plate geometry attached to the top plate and the nanoscale displacement applied at the 13 piezo points. The analysis presented in Figure 3 shows that the displacement in the wells of the cultureware increases with increasing frequency. This is likely due to the fundamental resonance of the top plate at 8.23 kHz, with the displacement amplitude starting to greatly increase as the resonance is approached. Even though the response is not as flat as anticipated, the bioreactor can be easily configured for 30 nm displacements across a 5 kHz frequency band with some moderate equalisation to the piezo drive signal. An average displacement of 28 ±2nm at 1 kHz is calculated at the bottom surface in each well. As expected, this is slightly less than the 30 nm displacement applied to the piezo points due to marginal losses during transmission of vibration from the piezo through the top plate to the cultureware.

**Figure 3.**
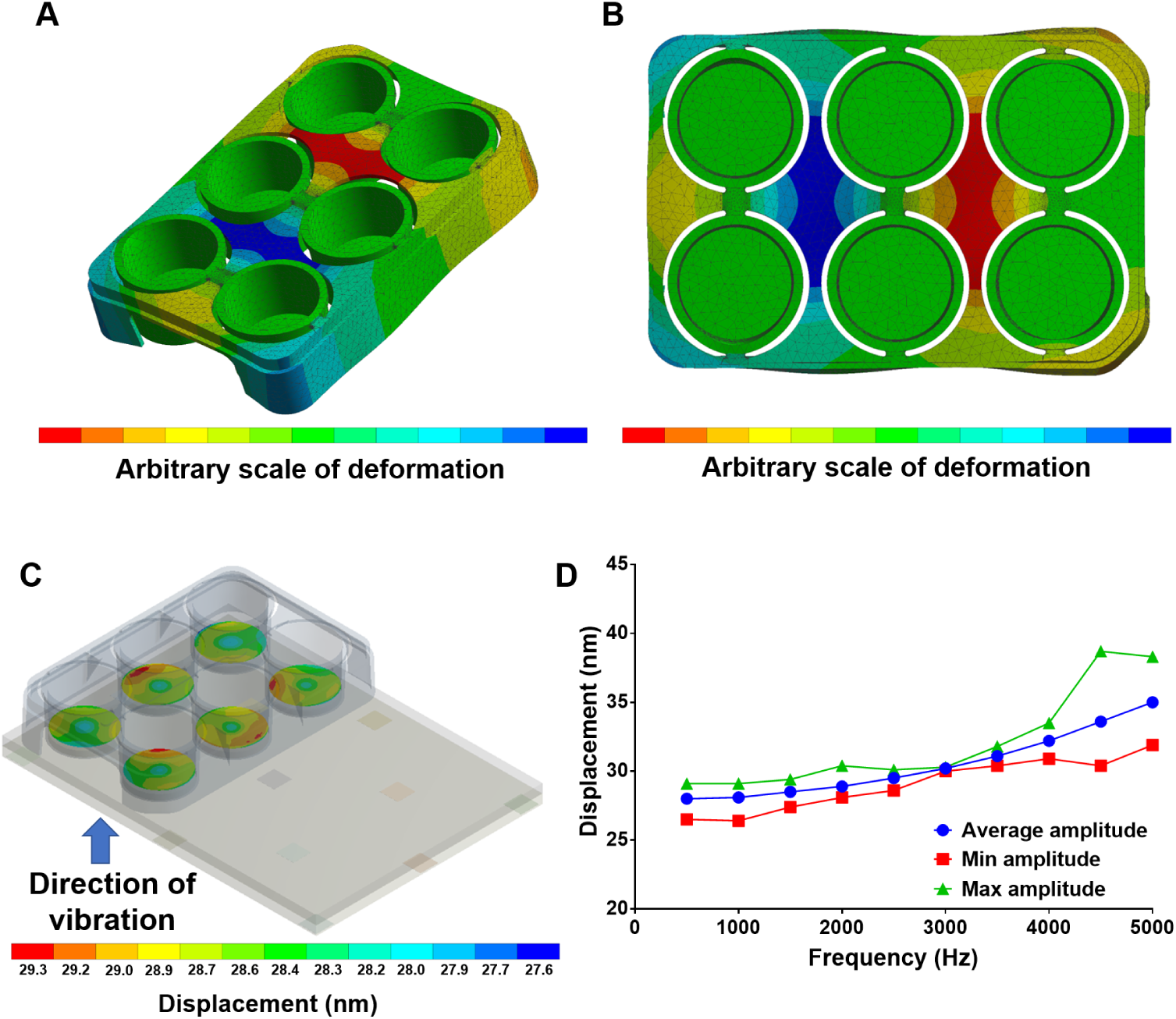
(A) and (B) show the mechanical response of the PP 6-well plate at its 12th mode (1037 Hz). Even though there are resonant modes in the bioreactor operating window, the resulting large scale displacement is confined to the frame and the wells themselves are unaffected. (C) Harmonic response analysis of combined 6-well plate and bioreactor top plate. (D) Frequency response of combined 6-well plate and bioreactor top plate component with fixed 30 nm displacement applied to the piezo bonding points.

### Other bioreactor vibration plate design refinements

The design of the bioreactor has been improved significantly over that presented in previous work^23^. The top plate of the bioreactor has been recessed into the base block to offer some protection from lateral mechanical shock which could risk damaging, or breaking, the epoxy bond between the piezos or the two halves of the top plate. Handling and portability of the bioreactor has been improved by providing handles and reducing the overall weight (with no noticeable effect on bioreactor dynamics). Drainage channels have been added to the base plate to allow cleaning fluids, and condensation, to drain easily – key considerations when planning for future GMP clean room use. Finally, stainless steel is now used for the top plate to prevent rusting in humid incubators, a known issue of the bioreactor used in the work by Tsimbouri *et al.*^23^. The bioreactor with PP cultureware is shown in Figure 4.

**Figure 4.**
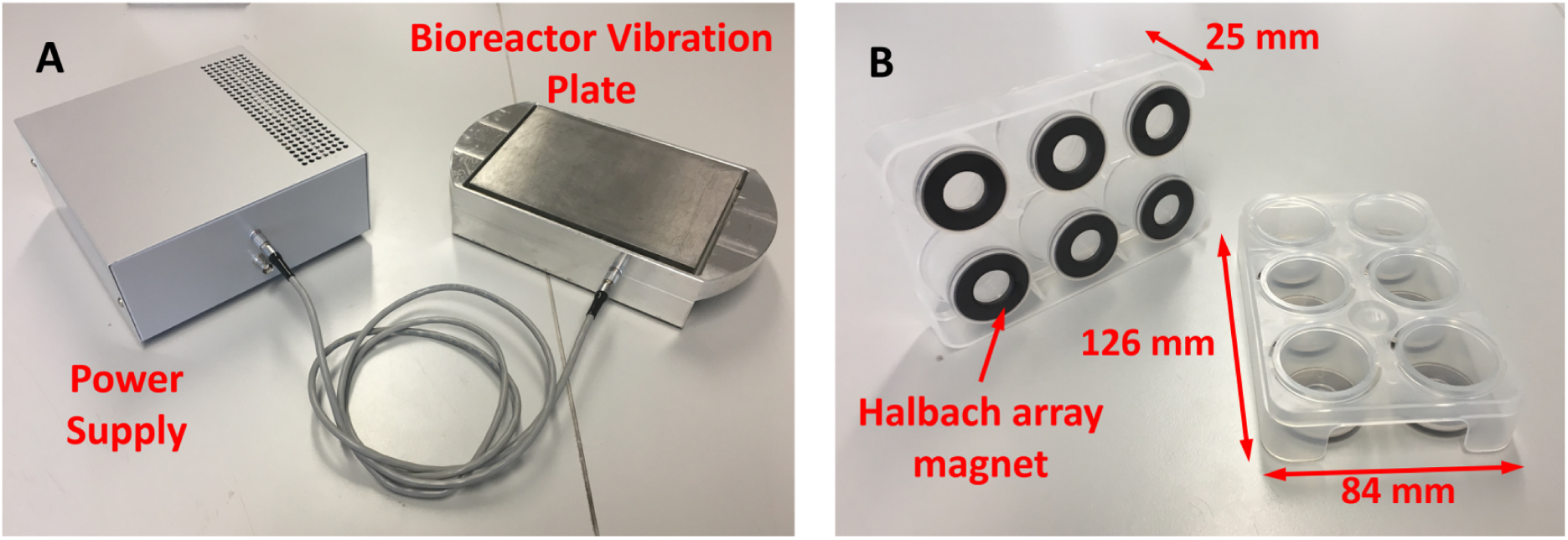
Bioreactor vibration plate with injection moulded PP 6-well cultureware. (A) The improved version of the bioreactor has a lighter base, carrying handles and a recessed top plate, along with a power supply designed to output a sine wave of 1 kHz and 30 nm displacement amplitude. (B) Injection moulded PP cultureware with incorporated halbach ferrite ring magnets in the base of each well.

### Power supply

A signal generator and a high-end audio amplifier had been previously used^23^ to supply a sine wave signal to the piezo array, requiring the user to have considerable knowledge of the equipment in order to achieve the desired displacements needed for their experiment. This has led to the development of a turnkey solution that allows users to simply connect the supply to the bioreactor and turn it on to get an average displacement of 30 nm at 1 kHz.

An AD9833 low power, programmable waveform generator (Analog Devices, Massachusetts, USA) produces the required sine wave with the output frequency and phase programmed using an ATMega328 microcontroller (Atmel, California, USA), allowing easy tuning. Appropriate filtering is implemented at the output of the AD9833 to significantly reduce higher frequency components, created during the digital synthesis of the primary signal^33^, using a seventh order LC elliptical reconstruction filter, ensuring that the 1 kHz sine wave drive signal is as pure as possible. A power spectrum of the pre and post filtered signal is shown in Figure 5. The filter is designed to have a cut-off frequency around 150 kHz and reduces the harmonics to a level where they do not have any observable effect on the bioreactor top plate displacement (based on sub-nm resolution measurements), therefore, the electrical harmonics are below the mechanical background of the system. The filtered AD9833 signal is inputted into an AC coupled op-amp, using an OPA37 ultra low noise precision amplifier (Texas Instruments, Dallas, USA), with variable gain to give fine adjustment of the amplitude of the sine wave before the main amplification stage using two TDA7293 class AB audio amplifiers (STMicroelectronics, Geneva, Switzerland) in a parallel configuration. These devices are primarily used for commercial HiFis; however, their low distortion and low noise characteristics, along with thermal shutdown and clip detection functionality, make them an excellent choice for this application. The parallel circuit configuration is based on the example circuit provided by STMicroelectronics in the TDA7293 datasheet^34^.

**Figure 5.**
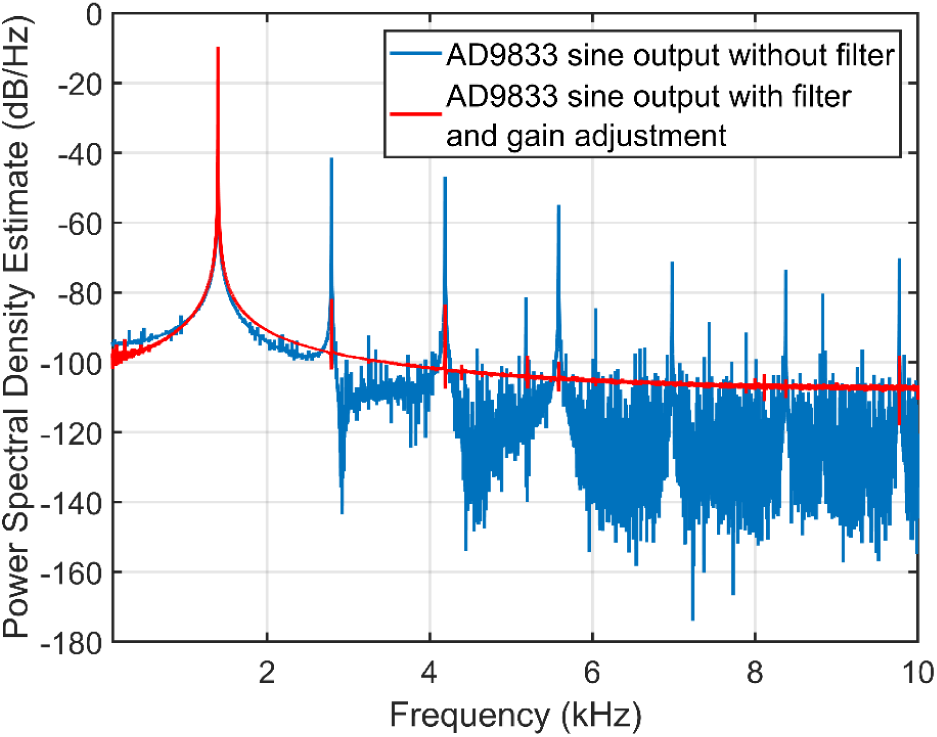
Estimate of power spectral density from AD9833 output pre/post filtering and gain adjustment The power spectral density of the output signal from the AD9833, after filtering and gain adjustment from OPA37 (red), shows a significant reduction in the harmonics of the generated drive signal. Even though some harmonics remain, they are well below the interferometer background noise level. Note that the frequency of the drive signal shown in this plot is 1.4 kHz.

### Interferometer characterization of bioreactor and cultureware

Verification of the FEA modelling and calibration of the bioreactor was carried out using a laser interferometer (Model SP-S SIOS Meßtechnik GmbH, Ilmenau, Germany). The interferometer is able to determine nanoscale changes in displacement from changes in the optical interference signal measured as laser light from the interferometer’s He-Ne source reflects off the target surface back to the photodiode detector, as illustrated in Figure 6. The displacement amplitude of the top plate was measured by reflecting the interferometer laser off a magnet (with prismatic tape attached, ensuring as much laser light as possible is reflected) on the top plate back to the receiver. The measurements of the top plate showed an average motion of 27 nm with a standard deviation of 33% across thirteen measurement points, roughly corresponding to the position of the piezos under the top plate. To measure the wells of the PP cultureware while magnetically attached to the bioreactor top plate, prismatic reflective tape was bonded to the bottom surface of each well. The average inter-well displacement amplitude measured was 25.4 nm with a standard deviation of 17.4%. The slight reduction in displacement amplitude is expected due to the additional layers between the vibration source and measurement site. In each well five points were measured to assess the uniformity of the displacement within a single well. Four of the six wells had standard deviations of less than 10% while the other two were slightly over 10%, with the largest standard deviation being 12.1%.

**Figure 6.**
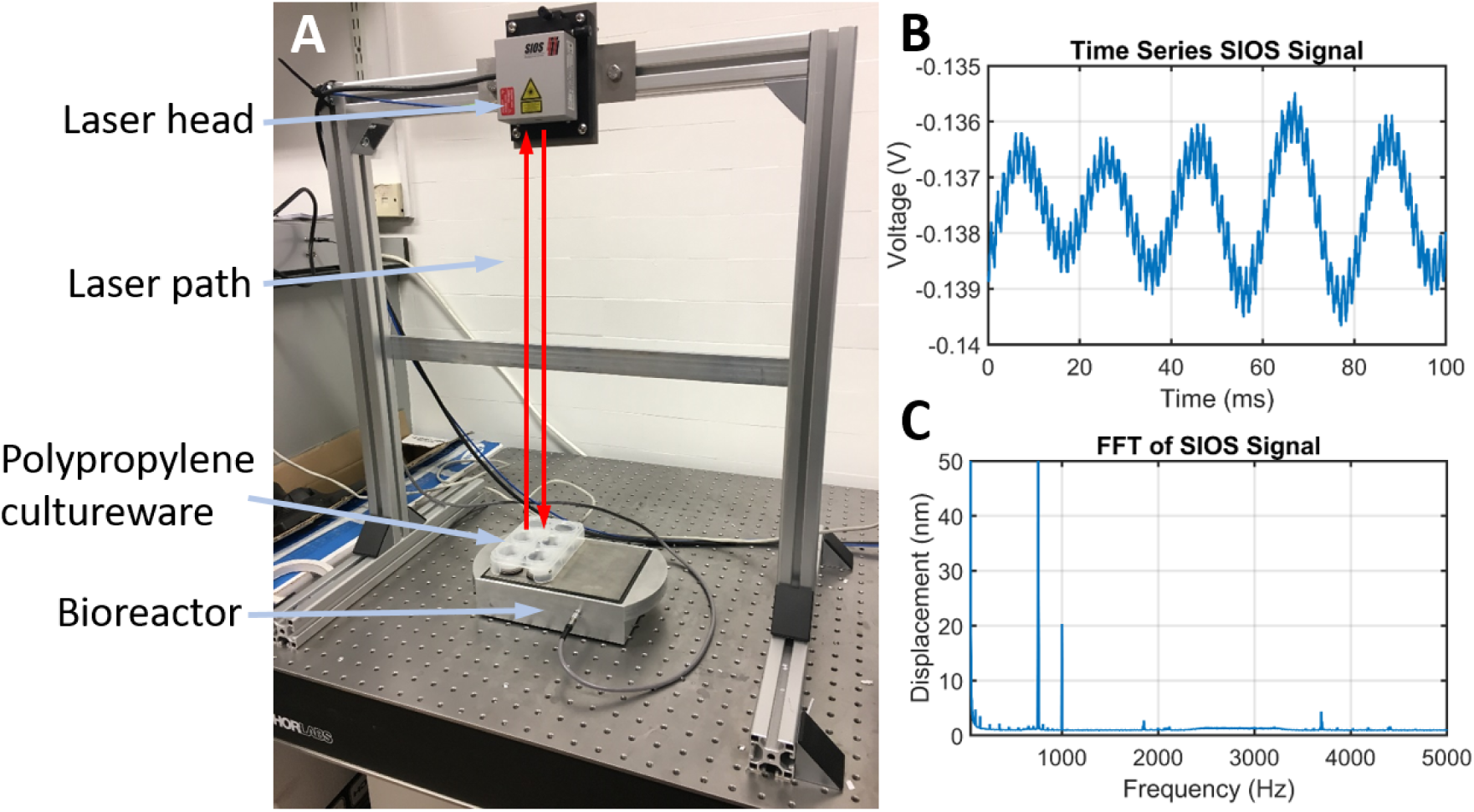
Interferometer measurement setup and output signal. (A) To measure nanoscale displacements the interferometer emits a laser beam from the laser head which is reflected back to the photodetector (also within the laser head) off the object being measured. Analysis of the optical interference pattern produced allows the displacement to be obtained. (B) Example of the time series data measured by the interferometer. (C) Example of an FFT analysis on the time series data. The 1 kHz peak of the bioreactor is clearly seen and there is also a large peak at 750 Hz, however, this signal is produced by the reference mirror of the laser of the interferometer which is constantly excited at a fixed frequency in order to obtain the control signals. Notice that the electronic harmonics discussed in the previous section have no effect at 2, 3, 4 and 5 kHz.

A similar methodology was used to compare the custom 6-well plate with other standard multi-well plates and tissue flasks coupled to the bioreactor via sheets of rubber magnet (also halbach arrays) bonded to the bottom surface of the cultureware. The data presented in Table 1 shows that the displacement amplitude is consistent over other multi-well culture plates and tissue flasks.

**Table 1.** Measurement of nanoscale displacements on custom PP 6-well plate and other standard PS tissue cultureware

| Plate Type | Average Displacement (nm) | Standard Deviation (nm) | Standard Deviation (%) |
| --- | --- | --- | --- |
| Custom 6-well plate | 25.4 | 4.4 | 17.4 |
| 6-well plate | 28.1 | 3.9 | 13.8 |
| 12-well plate | 26.4 | 3.3 | 12.3 |
| 24-well plate | 26.5 | 4.1 | 15.4 |
| T75 tissue flask | 25.1 | 3.2 | 12.9 |

### Estimation of accelerative force applied to cells

With precise values measured for the vibration amplitude, *A*_0_, along with the known frequency, *f*, the peak acceleration of the system can be quantified as *A*_0_(2*π f*)^2^. Using the peak acceleration and the mass of liquid media directly above one cell (cell surface area multiplied by media height and by media density), it is possible to estimate the force exerted on each cell using Newton’s second law. For the operating conditions of the system at 1 kHz and 30 nm amplitude, the acceleration experienced by the cells is approximately 0.12 *g*. Theoretically, for a constant amplitude of 30 nm the bioreactor should be able to apply accelerations from 0 *g* to 3 *g* in the range of 0 to 5 kHz. Being able to accurately define the forces the cells are experiencing allows our stimuli to be placed in context with other vibration and centrifugation studies, enabling the comparison of osteogenesis between waveform frequencies, amplitudes and accelerations.

### AFM measurements of collagen gel during nanovibration

The force calculation discussed in the previous section assumes that the cells are receiving a periodic (compressive/tensional) accelerative force during vibration which is acting on their membrane and cytoskeleton. Alternatively, the effect could be related to environmental stiffness, *i.e.* vibration could be causing strain hardening of the viscoelastic collagen gel, allowing cells to increase intra-cellular tension leading to downstream effects such as osteogenesis. Environmental stiffness is a factor that studies have shown impacts stem cell differentiation, with high elastic collagen gels in the 40 kPa range inducing osteogenesis in MSCs^35–39^.

To examine this possibility, AFM was used to determine any change in stiffness in a collagen gel whilst being nanovibrated. 2mg*/*ml collagen gel was measured with an aqueous layer of phosphate buffered saline above the gel surface, replicating the gel used in previous cell studies^23^. A tipless, silicon cantilever (Arrow TL1, Nanoworld) was used with an added 20 μm silicon bead to produce force-distance curves with a force setpoint of 3.0 nN. Due to the spatial limitation of the AFM system (JPK Instruments, Berlin, Germany) a single piezo actuator (PI model P-010.00H, Karlsruhe, Germany) was used to apply vibration to a Petri dish which was mounted under the AFM cantilever as shown in Figure 7. A signal generator (GW Instek, New Taipei City, Taiwan) was used to apply voltages of 0, 1, 5 and 10 Vpk-pk to the actuator at 1 kHz. A voltage signal of 20 Vpk-pk was attempted, however the AFM cantilever was unable to approach the collagen surface due to the increased motion. It was not possible to measure the precise vibration amplitude in situ but it is expected that this arrangement will deliver vibration amplitudes in the order of 9-14 nm at 10 Vpk-pk based on previous calibrations^21^.

**Figure 7.**
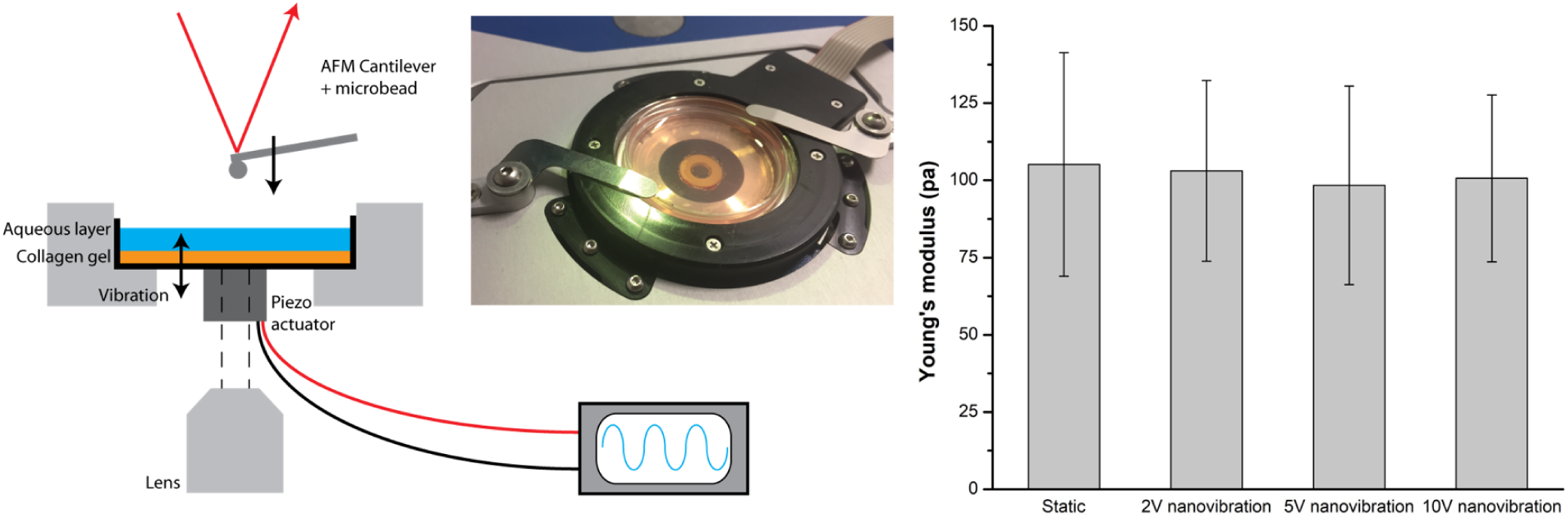
AFM measurement of collagen gel during nanovibration. (A) A vibrating piezo actuator was attached to a Petri dish containing collagen gel with an AFM cantilever being used to measure any changes in gel stiffness during vibration. (B) Young’s modulus was assessed via AFM for collagen samples which were nanovibrated at three different piezo amplitudes. Data are mean *±* SD (n > 30).

Vibration of the piezo introduced a periodic noise to the force-distance curves both during the cantilever approach (*i.e.* when in the aqueous layer) and when in contact with the collagen gel. This suggests that vibration is being transmitted through both the gel and the fluid above the gel supporting previous assumptions that the fluid acts as a rigid body when transmitting the accelerative force. In addition, the noise had a periodic component of 940 Hz which was estimated using the cantilever distance and extend speed (2 μm*/*s), although limited in accuracy by the sampling rate of 2048 Hz. This appears to be a direct measurement of the 1 kHz vibration signal. The amplitude of the noise was also found to increase in line with the vibration amplitude (see Supplimentary Fig. 2) with 10 Vpk-pk producing a peak force of 371 ± 11 pN when in contact with the gel (n = 6). Since we know the surface area of the cantilever tip (10 μm radius) we can estimate that a cuboidal cell of 140 × 100 μm would receive a peak force of 17 nN due to this vibration (9 nm, 1 kHz). It is worth noting that this measurement is largely consistent with our previously calculated estimates in the 10s of nN for vibration of 30 nm at 1 kHz^20–22,40^.

Although noise increased the residual RMS of the Hertz model curve fitting it was still possible to collect data on the stiffness of the gel during nanovibration. No significant effects of strain hardening were observed within the gel and the Young’s modulus remained in the region of 100 Pa (Figure 7) which is consistent with the values of the soft collagen gels used in work by Tsimbouri *et al.*^23^. n > 30 stiffness measurements were included in calculations of Young’s modulus (taken over two collagen samples). Again, we note that osteogenesis is only promoted in static gels with stiffnesses in the kPa range^35–39^.

### Biological validation

As stated, previous work^20, 23^ has shown moderate up-regulation of genes associated with a switch towards an osteogenic lineage. However, protein expression is a more definitive measure of this switch. Nanovibrational stimulation was applied to MSCs for three weeks and compared against non-stimulated MSCs. As shown in Figure 8, nanovibrational stimulation of MSCs for three weeks resulted in a significant increase (p < 0.05) in the known osteogenic-related protein expression of Runt-related transcription factor 2 (RUNX2), osterix (OSX), osteopontin (OPN), osteocalcin (OCN) and ALP, compared to the non-stimulated control. These proteins are markers of osteogenic differentiation, signifying that the new design of bioreactor is capable of stimulating MSC osteogenesis in line with previously reported results. Detailed information about the cell culture and In-Cell Western processing can be found in the Methods section.

**Figure 8.**
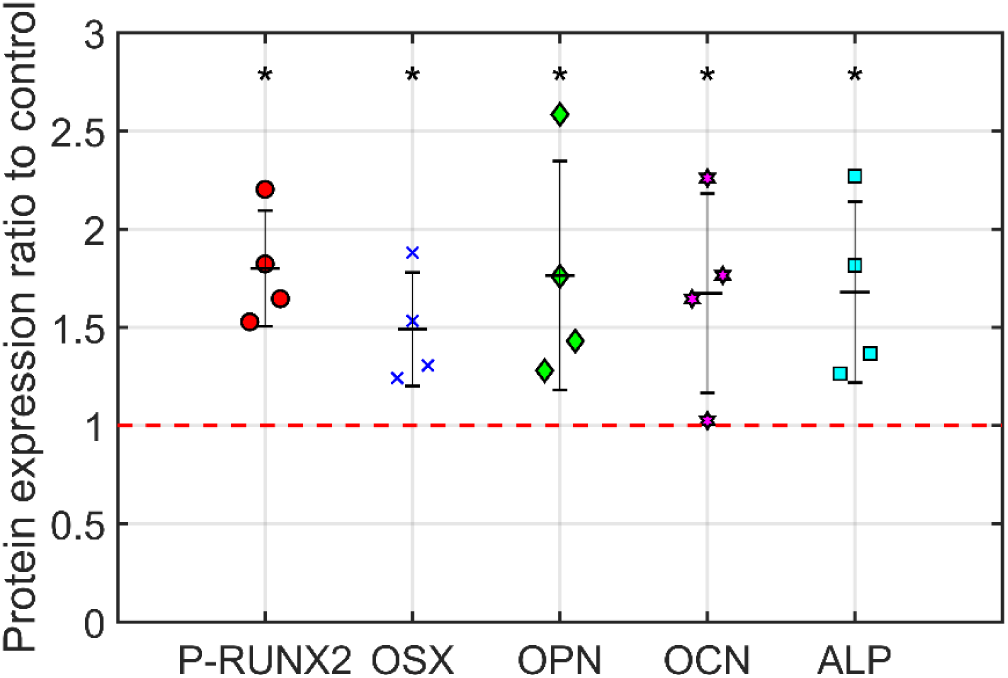
Protein expression at three weeks of nanovibrational stimulation. Cells were stimulated for three weeks at a frequency of 1 kHz and 30 nm displacement alongside a non-stimulated control. Protein expression for Runx2, OSX, OPN, OCN and ALP, was measured relative to static controls and quantified using the LI-COR Odyssey system (LI-COR, Nebraska, USA). Statistically significantly higher expression was observed in the mechanically stimulated samples when compared to the non-stimulated control (indicated by red dashed line). Data are mean ± standard deviation, n = 4, stats Mann Whitney U-test * p < 0.05.

## Discussion

The bioreactor system described in this article works well on the scale of a research project, or small clinical trial, that requires mechanical stimulation with an amplitude of approximately 30 nm at a frequency of 1 kHz. The design and development of the system has been approached with a GMP ethos to ensure that all bioreactors built are consistent in their vibrational output, while also being cost effective, scalable and simple to use so that the system can be widely used by cell therapy manufacturers and researchers with reproducible results. The bioreactor system has already been sent to a number of research labs, running successful experiments with minimal training from the authors here and using a variety of cultureware formats, as validated in this paper. Infact, the first clinical evaluation of surgical bone graft using mechanically stimulated MSCs, created with this bioreactor platform, and a 3D polycaprolactone (PCL) construct coated in layers of poly(ethyl acrylate) (PEA), fibronectin (FN) and growth factor BMP-2^41–43^ is scheduled for 2020/21, funded by the landmine charity Find A Better Way^44^. As well as the cost-effective production of the system, there is also the possibility of substantial cost savings to be made by healthcare organisations when using the bioreactor to produce cellular bone graft material for surgery in orthopaedic patients. Conceptually, allogenic bone cells present significant ‘off-the-shelf’ benefits over the current orthopaedic gold standard - harvesting and transplanting autograft from the patient’s own iliac crest by minimising risk associated with opening a second surgical site.

Due to the growing interest in mechanotransduction as a method to control cell behaviour and stem cell differentiation, the authors are developing a new power supply unit that can deliver a broader range of user-defined frequencies (1 Hz to 5 kHz) and amplitudes (up to 100 nm). If cellular nanoscale mechanotransduction/ mechanosensing has indeed developed early within evolutionary history, as suggested by the bacterial studies conducted^45^, then further exploitation of the bioreactor technology is highly likely. Thus, an entirely new field of mechanobiology may be opened. Furthermore, the bioreactor could have as yet unforeseen potential when used to stimulate other cell types within its measured frequency and acceleration ranges. Studies involving suppression of certain cancer cell lines and pathogenic organisms (bacteria) are currently being investigated^45^.

Future work will also be aimed at developing a system that can provide mechanical stimulation to a much higher quantity of cells. At the point where large scale clinical trials are being conducted, we may need to consider a different approach to the system to increase the number of cells produced to the region of 100s of millions instead of 100s of thousands. We would also like to improve upon the FEA modelling by exploring different interferometric and imaging techniques.

## Methods

### FEA analysis

3D computer aided drawing (CAD) models of the bioreactor top plate and PP cultureware were developed in PTC Creo parametric 3.0 (PTC, USA) and imported into ANSYS workbench 17.1 (ANSYS, Canonsburg, PA) for FEA. The two analysis types used were Modal and Harmonic Response. A Modal analysis produces estimations of the internal resonant modes of the model and the Harmonic Response analysis shows deformation of components due to nanoscale vibration. All geometries were meshed using the patch conforming tetrahedron method, with the following sizing options for mesh resolution: Relevance Centre – Fine, Smoothing – high, Transition – slow, and Span Angle Centre – fine. It was assumed that, as the ferrite magnets are encapsulated in the culture plate, there is a rigid bond between the magnets and the contacting PP surfaces. The magnetic face of the magnets is bonded to the magnetic surface of the top plate for this model.

Modal analysis was performed to identify the first 12 modes for the top plate and the custom cultureware. Fixed supports were applied to the underside of the epoxy of the top plate and 6 magnets for custom cultureware. For the harmonic response analysis displacement of 30 nm was applied to the underside of the 13 epoxy layers for the top plate and the 6 magnets for custom cultureware. An inertial acceleration of 9.806 m*/*s^2^ was applied in the opposite direction of the piezo forces to simulate gravitational loading.

### Laser interferometry

Verification of the FEA modelling and calibration of the bioreactor was carried out using a laser interferometer (SIOS Meßtechnik GmbH, Ilmenau, Germany) as previously described^23^. A Fast Fourier Transform (FFT) of the time series measurement allows us to determine the displacement that is only created by the piezos at the chosen output frequency. Confirmatory data for the FEA was taken by first measuring the top plate without any cultureware and then by measuring the cultureware on the top plate. The top plate was measured by reflecting the interferometer laser off a magnet on the top plate. The magnet had prismatic reflective tape attached to it to ensure that as much laser light as possible is reflected back to the receiver. For the next stage of measurement, a 6-well plate was magnetically attached to the top plate of the bioreactor. As with the previous stage of measurements, reflective tape was attached to the bottom of each well to return the interferometer laser back to the receiver.

### AFM measurement

2 mg*/*ml collagen gel was measured with an aqueous layer of phosphate buffered saline above the gel surface. A tipless, silicon cantilever (Arrow TL1, Nanoworld) was used with an added 20 μm silicon bead to produce force-distance curves with the force setpoint set to 3.0 nN. AFM data was collected in force spectroscopy mode using Nanowizard software (JPK Instruments, Berlin, Germany) and JPK processing software was used to fit force curves with the Hertz model to determine the Young’s modulus.

### Mould design and flow analysis

The mould tool was manufactured by CGP Engineering Limited with mould base components made of 1.1730 Carbon Tool Steel (C=0.45, Si=0.30, Mn=0.70), materials supplied by Milacron. The mould was manufactured using mould making computer numerical control (CNC) machines. Draft walls, difficult to machine areas were machined with CNC driven electro-discharge machining process. Manufactured mould tool was mounted on a Demag NC4 Ergotech 100 – 310 injection moulding machine for initial prototypes. The following injection moulding parameters for the flow simulation were taken through initial prototyping and experimentation on the moulding machine: melt temperature of 205 ^*◦*^C, mould temperature of 15 ^*◦*^C, maximum injection pressure of 350 Bar, clamping pressure of 500 KN, cooling cycle time of 20 seconds with overall moulding cycle time of 55 seconds for each injection cycle. Culture-ware design was revised through this analysis in order to produce defect free product. Details of flow simulation which informed the moulding process can be found in Supplementary Results.

### Water contact angle

WCA was measured on air plasma treated injection moulded culture plates using the sessile drop method. A Theta optical tensiometer was used (Biolin Scientific, Gothenburg, Sweden). For each condition and time-point, triplicate measurements were taken by placing 3 μl of Milli-Q water on the sample and the software used to fit the droplet curvature using the Young-Laplace equation. Samples were stored at room temperature during and between measurements.

For the cell culture primary bone marrow MSCs were sourced (Promocell, Heidelberg, Germany. Expanded MSCs were used at passages 1-3. The cells were maintained in basal media consisting of DMEM (Sigma Aldrich, St. Louis, USA) supplemented with 10% FBS (Sigma), 1% sodium pyruvate (11 mg/ml, Sigma), 1% Gibco MEM NEAA (amino acids, Thermo Fisher Scientific, Waltham, USA) and 2% antibiotics (6.74 U/mL penicillin-streptomycin, 0.2 μgμl^*−*1^ fungizone). Prior to every experiment, cells were trypsinised and counted with a haemocytometer. Cell concentration was typically 10,000 cells/ml. All cell culture was performed in an incubator at 37 ^*◦*^C with 5% CO2. The basal culture media was removed and replenished every 2-3 days.

### In-Cell Western

MCSs were fixed in a 10% formalin sucrose solution for 15 min at 37 ^*◦*^C, permeabilised in permeabilisation buffer pH 7.2 (30 mM sucrose, 50 mM NaCl, 7 mM MgCL2 (hexahydrate), 20 mM HEPES, 0.5% Triton X-100 in 1 x PBS), and blocked in 1% (w/v) milk/PBS for 1.5 hr at 37 ^*◦*^C on a plate shaker. MSCs were stained using primary antibodies 1:200 anti-PRUNX2 (rabbit, pS465 Abgent), ALP (sc-166261, Santa Cruz Biotechnology), OCN (sc-73464, Santa Cruz Biotechnology), OPN (sc-21742, Santa Cruz Biotechnology,) OSX (sc-22533, Santa Cruz Biotechnology), GapDH (sc-32233, Santa Cruz Biotechnology) and CD90 (13-0909-82, eBiosciences). The primary antibody incubation was carried out at 37 ^*◦*^C for 2.5 hr, following 5 washes in PBS/0.1% Tween. GAPDH and CellTag 700 stain (Li-COR, cat: 926-41090, Lincoln, USA) were used as the reference controls. CellTag or rabbit anti-GAPDH primary antibody was diluted at 1:2000. The Li-COR secondary antibodies (anti-rabbit, cat: 926-32211; anti-mouse, cat: 926-32210) were used at a dilution of 1:2000. For GAPDH, the secondary antibody was diluted at a dilution of 1:5000 with a 680 nm infrared dye. The secondary antibodies were incubated at room temperature on a shaker for 1.5 hr, followed by 5x 5 min washes in PBS/0.2% Tween. The quantitative spectroscopic analysis was carried out using the LI-COR Odyssey Sa (LI-COR, Nebraska, USA).

## Acknowledgements

The authors would like to thank: Iain Martin, Habib Nikukar, Keith Robertson, Gilad Tiefenbrun and Ivor Tiefenbrun for their advice, Gerry O’Hare and Ross Simpson at the University of the West of Scotland for their help etching and assembling the power supply PCBs, Jim Orr and his workshop team at the University of the West of Scotland for machining the bioreactor parts and for helpful discussions on their design, CGP Engineering Limited for manufacturing the mould and carrying out the initial production run of cultureware, Milacron for advice and useful discussion on mould design and materials, and the GEO and LIGO Scientific Collaboration for their interest.

Funding and financial support from STFC (ST/N005406/2, ST/L502509/1), BBSRC (BB/N012690/1, BB/P00220X/1), EPSRC (EP/N013905/1, EP/P001114/1), Find A Better Way, SUPA, the Royal Society (RS), the Royal Society of Edinburgh (RSE), NHS Greater Glasgow & Clyde, Linn Products Ltd, the University of the West of Scotland, University of Glasgow and University of Strathclyde are gratefully acknowledged.

## Author contributions statement

P.C. designed and constructed the PSU, performed laser interferometry and drafted the manuscript. P.G.C. performed AFM measurements, cell adhesion/WCA experiments and drafted the manuscript. S.N.R. performed ANSYS modelling of bioreactor and cultureware and drafted the manuscript. K.C. and P.V. assisted in design of cultureware and injection mould tooling. J.H. helped with initial design considerations for the system and advised on the precision measurement techniques used. M.P.T. carried out MSC experiments with the bioreactor and performed protein expression assay. M.S.S., M.J.D. and S.R. lead this research in their respective departments and helped draft the manuscript. All authors read and approved final manuscript.

## Availability of data and material

The datasets used and/or analysed during the current study are available from the corresponding author on reasonable request.

## Additional information

**Supplementary information** accompanies this paper

