## Supplementary Information for "Design, construction and characterisation of a novel nanovibrational bioreactor and cultureware for osteogenesis"

### Supplementary Results

#### Custom cultureware design

**Supplementary Fig. 1A** presents an exploded view of the developed mould, illustrating major components of the mould interface and ejection system. Six sleeve ejectors in the ejector mechanism are guided on magnet holding pins, ejecting moulded cultureware. The design was evaluated for mouldability using Moldex3D (Creo Mold Analysis Extension 4.0) mould flow simulation software. Simulating mould flow through the mould cavity provides potential void areas, mould filling confidence and fill progression. Injection pressure drop during polymer filling phase and many potential plastic defects such as sink marks and weld lines were also taken into account during simulation in order to create a defect free product (see **Supplementary** **Fig. 1B**). The filling time for each cycle was determined to be 3.65 seconds with a cooling cycle time of 20 seconds and overall moulding cycle time of 55 seconds for each injection cycle.

**Fill Progression**

**25% Fill**

**50% Fill**

**100% Fill**

**Av**

Moving side of

mould tool

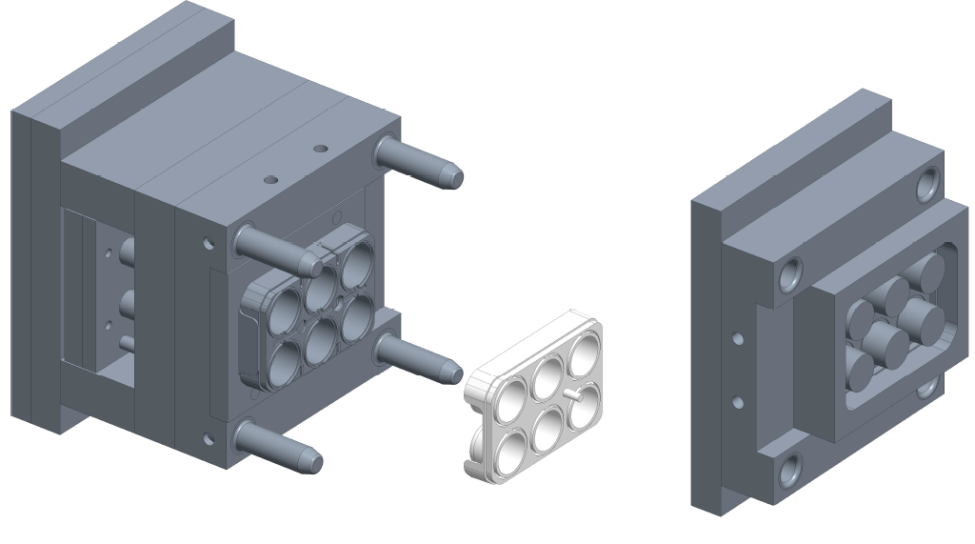

Moulded culture-ware

Ring magnets

Ejection system

Injection side of

mould tool

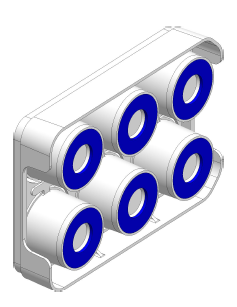

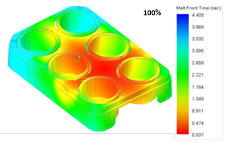

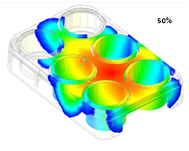

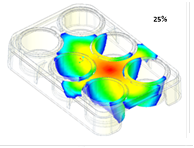

**Bv**

**Figure 1: Injection mould tool design and mould fill analysis**

(A) An exploded view of the mould tool with culture plate is shown to illustrate the major components of the mould interface and ejection system. (B) shows the mould fill analysis which estimates that the part cavity in the tool should take 3.65 seconds to completely fill giving a defect free part.

For the injection moulding process polystyrene (PS) was initially chosen as the material for the cultureware as it is commonly used for off the shelf brands. However, during injection mould production trials it was found that the 6-well plate would crack easily when the ejector pins of the mould tool pushed the plate out of the tool at the end of the moulding process. Due to the floating well design of the plate, there was a reduced number of ejection sites in comparison to standard 6-well cultureware plates. It was assumed that this lead to increased force being exerted on the wells, cracking the PS part. This issue necessitated a change of material, polypropylene (PP) was chosen as it is still amenable to surface modification, has increased flexibility compared to PS and is also used in standard cultureware. Trial runs with PP demonstrated that the cracking problem had been eliminated.

#### Custom cultureware modal analysis

FEA was performed on the 6-well plate design with a modal analysis revealing that the cultureware has a fundamental mode at 379 Hz and some modes close to 1 kHz (for table of modes 1 to 12 see **Supplementary Table S1**).

**Supplementary Table S1**: First 12 resonant modes of PP cultureware determined from ANSYS modal analysis

| **Mode** | 1 | 2 | 3 | 4 | 5 | 6 | 7 | 8 | 9 | 10 | 11 | 12 |
| --- | --- | --- | --- | --- | --- | --- | --- | --- | --- | --- | --- | --- |
| **Frequency (Hz)** | 379 | 381 | 500 | 547 | 584 | 660 | 772 | 840 | 903 | 933 | 961 | 1037 |

#### AFM measurements of collagen gel during nanovibration

Vibration of the piezo in the AFM set-up introduced a periodic noise to the force-distance curves both during the cantilever approach (*i.e.* when in the aqueous layer) and when in contact with the collagen gel. The amplitude of the noise increased with vibration amplitude, as shown in Supplementary Fig. 2, however, this did not greatly affect the estimates of the Young’s modulus from the force-distance curves.

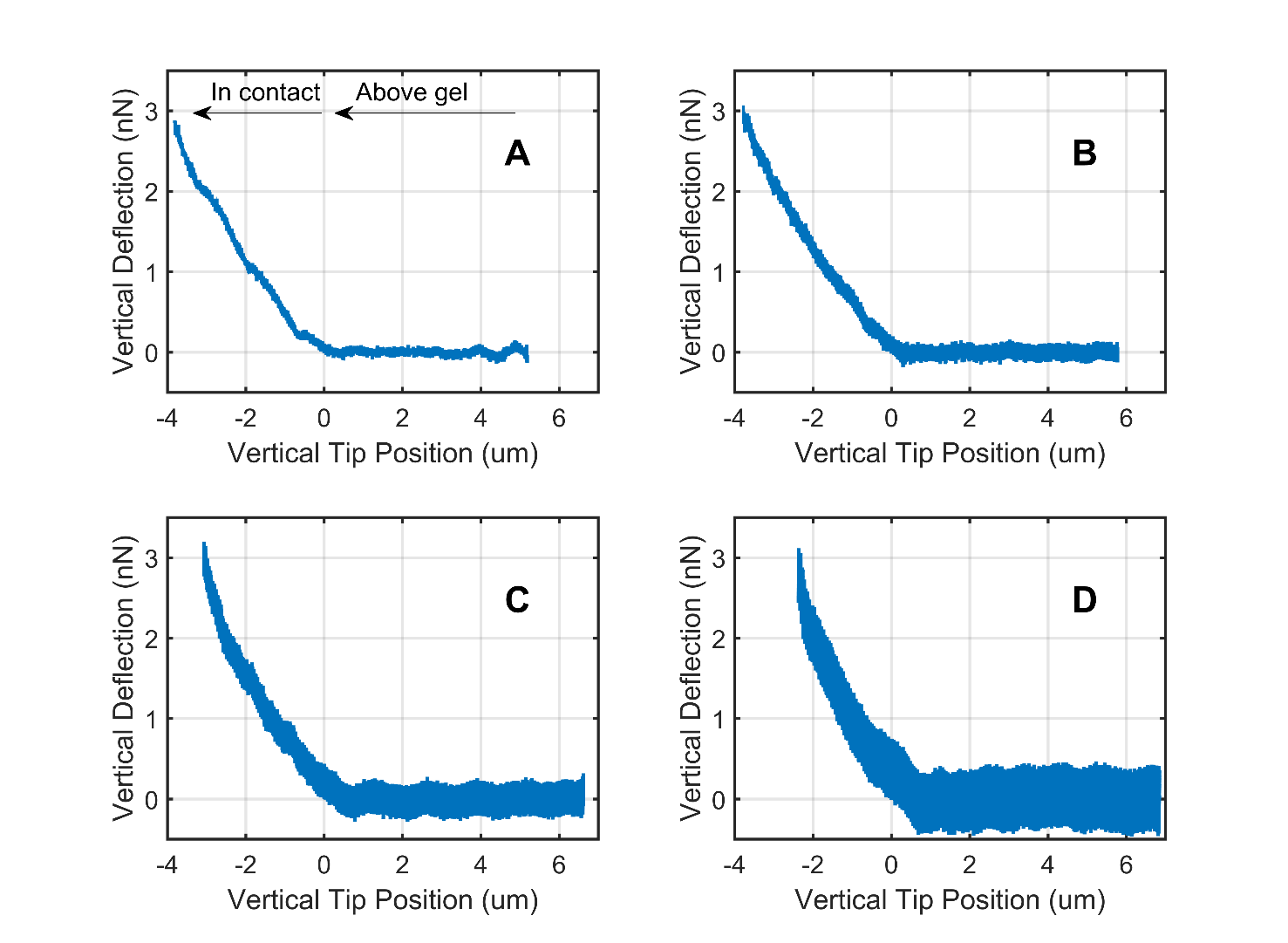

**Figure 2: AFM force-distance curves of nanokicked collagen**

Example force-distance curves of collagen gels measured via AFM during nanovibration for A) No vibration, B) 2 V_pk-pk_ vibration, C) 5 V_pk-pk_ vibration, D) 10 V_pk-pk_ vibration. The force applied by the nanovibration is visible as amplitude dependant noise.

### **Supplementary Method**

#### **Mould design and flow analysis**

Cero parametric 3.0 (PTC, USA) Product Lifecycle Management (PLM) system was utilized for i) Capturing all product requirements ii) Culture-ware design and iii) Mould design. Culture plate design was evaluated for mouldability using Moldex3D (Creo Mold Analysis Extension 4.0) mould flow simulation software. Polypropylene grade of Eltex® MED 240-MS23 (supplied by INEOS O&P Europe) was used for initial prototypes and required material parameters needed for the mould flow analysis were obtained from the data sheet [1]. The following injection moulding parameters for the flow simulation were taken through initial prototyping and experimentation on the moulding machine: melt temperature of 205°C, mould temperature of 15°C, maximum injection pressure of 350 Bar, clamping pressure of 500KN, cooling cycle time of 20 seconds with overall moulding cycle time of 55 seconds for each injection cycle. Culture-ware design was revised through this analysis in order to produce defect free product. These parameters are then fed forward to Injection molding machine for producing physical prototypes.
